# Low maternal plasma apelin in the second trimester is associated with adverse pregnancy outcome in severe, early-onset fetal growth restriction

**DOI:** 10.1101/2024.09.19.613895

**Authors:** N. Braun, K. Bethell, L. Chaloner, K. Maksym, R.N. Spencer, J.J. Maguire, A.P. Davenport, EVERREST consortium, A.L. David, O.R. Vaughan

**Affiliations:** Department of Maternal and Fetal Medicine, Elizabeth Garrett Anderson Institute for Women’s Health, University College London, London, United Kingdom, WC1E6HX; Leeds Institute of Cardiovascular and Metabolic Medicine, University of Leeds, UK, LS2 9NL; Experimental Medicine and Immunotherapeutics, University of Cambridge, Addenbrooke’s Hospital, Cambridge, United Kingdom, CB2 0QQ; National Institute for Health and Care Research University College London Hospitals Biomedical Research Centre, London, United Kingdom, W1T 7DN

**Keywords:** apelin receptors, intrauterine growth restriction, ELA peptide, human, premature birth, stillbirth

## Abstract

**Context:** Fetal growth restriction increases adverse pregnancy outcomes such as preterm birth and intrauterine fetal death. Apelin is a secreted peptide expressed in placental syncytiotrophoblast and downregulated in fetal growth restriction.

**Objectives:** We tested the hypothesis that adverse pregnancy outcome is associated with low maternal plasma apelin at diagnosis of early-onset fetal growth restriction.

**Methods:** Plasma samples and fetomaternal blood flow Doppler velocimetry measurements were obtained from pregnant women (n=59) at diagnosis of early-onset fetal growth restriction in the second trimester. Plasma apelin was determined by ELISA and pregnancy outcome was recorded. Placental gene expression was analysed after birth by qRT-PCR, compared to term placentas from women with late-onset fetal growth restriction or with appropriate-for-gestational age infants.

**Results:** At diagnosis of early-onset fetal growth restriction, plasma apelin concentration was significantly lower in women who delivered extremely preterm (<28 weeks gestation) or had an intrauterine fetal death, compared to women who had a livebirth≥28 weeks (P<0.05). Plasma apelin correlated directly with uterine artery volume flow rate and inversely with pulsatility index. Placental gene expression of apelin, but not the apelin receptor or elabela, was lower in women with early-onset fetal growth restriction delivering preterm than in appropriate-for-gestational-age, term control women.

**Conclusion:** Low maternal circulating apelin during the second trimester is associated with impaired uteroplacental perfusion and subsequent adverse pregnancy outcome in severe, early-onset fetal growth restriction. Placental apelin deficiency may contribute mechanistically to the pathogenesis of early-onset fetal growth restriction.

## INTRODUCTION

Fetal growth restriction (FGR) affects 5-10% of pregnancies and is characterized by suboptimal fetal growth [1]. FGR increases the risk of preterm birth and is the second leading cause of perinatal death [2]. A key obstacle to lowering mortality and morbidity is accurately diagnosing and managing FGR. Even in high-income countries, over 50% of FGR cases are undiagnosed during pregnancy [3, 4], while 70% of pregnancies ending in FGR-related intrauterine fetal death (IUFD) are not identified by sonography [5]. FGR is often associated with placental dysfunction, with reduced uteroplacental blood flow and defective nutrient transport across the syncytiotrophoblast epithelium. There are no approved treatments for FGR and the underpinning mechanisms remain poorly understood, preventing the development of improved screening tests and therapies.

Early-onset FGR occurs before 32 weeks gestation whereas late-onset FGR occurs after 32 weeks gestation. Both are diagnosed by ultrasound biometry measurement of small fetal size in combination with Doppler velocimetry measurement of abnormal uterine or umbilical artery resistance [6]. Early-onset FGR is strongly associated with pre-eclampsia, maternal vascular malperfusion, fetal hypoxia, preterm delivery and fetal demise [7–9]. Contrastingly, late-onset FGR is less commonly associated with pre-eclampsia, vascular abnormalities or fetal mortality but affects a larger proportion of pregnancies, is a common cause of stillbirth and carries a similar burden of poor long term neurodevelopmental and cardiometabolic outcomes in the neonate [7]. The distinct pathophysiological mechanisms contributing to early- and late-onset FGR have not been investigated.

The apelin system comprises the apelin receptor (APLNR) and two secreted peptides, apelin and elabela [10]. APLNR is a membrane-bound G protein-coupled receptor that is ∼50% homologous to the angiotensin II receptor [10]. Apelin and elabela are encoded by *APLN* and *APELA* genes, respectively *[10]*. Apelin is a powerful modulator of the cardiovascular, metabolic and reproductive systems, including acting as a cardiac inotrope, vasodilator and insulin sensitiser [10].

There is evidence to suggest that the apelin system plays an important role in placental function. Apelin is highly abundant in the placenta and both the ligand and APLNR are expressed in trophoblast cells, including the syncytiotrophoblast [11–15]. FGR and pre-eclampsia are associated with diminished placental apelin and APLNR abundance [13, 14, 16]. Maternal circulating apelin concentrations are also lower in women with FGR, when compared to uncomplicated pregnancies, and in pregnant rats that are experimentally undernourished to reduce fetal growth [14, 17]. Apelin promotes proliferation, invasion, hormone secretion, and nutrient transport in trophoblast cells *in vitro* and in rodent placentas *in vivo* [11, 17–20]. Apelin deficiency may therefore contribute mechanistically to placental dysfunction in FGR.

The aim of this study was to determine the association between maternal circulating apelin abundance in the 2^nd^ trimester, uteroplacental perfusion and pregnancy outcome, in women diagnosed with severe early-onset FGR. We hypothesised that adverse pregnancy outcome is associated with low maternal plasma apelin at diagnosis with FGR. We also measured placental expression of apelin system components separately in early-onset FGR compared to late-onset FGR and appropriate-for-gestational-age controls.

## METHODS

### Study participants and sample collection

Participants were recruited with written, informed consent under studies ethically approved by a UK National Health Service Research Ethics Committee (Stanmore, 13/LO/1254 or Hampstead, 15/LO/1488).

Pregnant women diagnosed with severe, early-onset FGR (n=59) were prospectively recruited at gestational ages between 20+0 and 26+6 weeks (+days) from four European tertiary referral centres (University College Hospital London, University Medical Centre Hamburg-Eppendorf, Maternal-Fetal Unit Hospital Clinic Barcelona, and Skane University Hospital Lund). Women with a singleton fetus and ultrasound estimated fetal weight (EFW) below 600 g and <3^rd^ centile for gestational age were eligible. Exclusion criteria included multiple pregnancy, maternal age <18 years, fetal structural or karyotypic abnormalities or maternal infection.

At recruitment, maternal venous blood was collected into EDTA-coated tubes then centrifuged to obtain plasma. Samples were aliquoted and stored at −80°C within 30 minutes of collection. Fetal biometry and uterine and umbilical arterial and venous Doppler velocimetry indices were also measured on the day of recruitment using a standardised ultrasound protocol. Estimated fetal weight and was calculated according to the Hadlock 3 equation[21] and the z-score computed as described [22]. Uterine artery and umbilical vein volume flow rates were determined from the averages of three measurements of blood flow velocity and vessel diameter, also as described [23]. Participants received standard care, in line with local guidelines, for management of early-onset FGR through pregnancy and delivery. Pregnancy outcome was categorised into one of two groups, based either on (1) fetal intrauterine death or extremely preterm delivery <28 weeks, or (2) livebirth ≥28 weeks.

Pregnant women with late-onset FGR (n=8) or carrying appropriate-for-gestational-age (AGA) control fetuses (n=15) were recruited at term when they delivered at University College Hospital London. Participants were eligible for the late-onset FGR group if the fetus was appropriately grown with EFW >10^th^ centile for gestational age at the mid-gestation anomaly scan but had an EFW <10^th^ centile after 32+0 weeks of gestation. Exclusion criteria were the same as for the early-onset FGR group.

For all study groups, if the fetus was liveborn, placental villous tissue was systematically sampled from two areas of the placental parenchyma, both midway between the umbilical cord insertion and margin. Samples were dissected free from the decidua and chorionic plate, preserved in RNALater and stored at −80°C.

### Plasma apelin measurement

Plasma samples were cleaned using an established solid-phase extraction method [24, 25]. Briefly, plasma was thawed on ice and 400 μl of each sample was transferred to a prechilled microcentrifuge tube (LoBind, Eppendorf) containing 100 μl of hydrochloric acid (2M HCl). Tubes were vortex-mixed and then centrifuged (20,000 rpm, 20 min, 4°C). The supernatant was loaded onto a 96-well Oasis Primed HLB μ-Elution plate. The elution plate was centrifuged (200 rpm, 5 min, 4°C), the wells washed with 200 μl of wash buffer (5% methanol, 1% acetic acid in H_2_O) and the plate centrifuged again (200 rpm, 5 min, 4°C). Samples were eluted into a clean 96-well Protein LoBind plate with 3x 50 μl of elution buffer (60% methanol, 10% acetic acid in H_2_O) and centrifuged (200 rpm, 5 min, 4°C). The eluate was then evaporated to dryness using a vacuum centrifuge (1200 rpm, 5 min, 4°C).

Dried down samples were resuspended in 120 μl of assay buffer and centrifuged to eliminate foam (1200 rpm, 5 min, 4°C). The samples were then added in duplicate to the pre-coated wells of a commercially available apelin ELISA kit and assayed according to manufacturer’s instructions (Phoenix Pharmaceuticals, Inc.; Catalog number: EK-057-23; Lot number: 610671). The minimum detectable concentration for this kit was 0.08 ng/ml, while the intra-assay and inter-assay variation was <10% and <15%, respectively. The detectable range was 0 to 100 ng/ml. The ELISA kit used in this study detected apelin-12 and had 100% cross-reactivity to apelin-13 and apelin-36. The final colour change reaction was measured at 450 nm using a Synergy HT microplate reader (Biotek, Vermont, USA). GraphPad Prism software version 9.5.1 for Windows (GraphPad, Software, USA) was used to create a sigmoidal curve of standard apelin-12 concentrations versus absorbance and interpolate the concentrations of the participant samples. The average concentration of the positive control was within the 0.2-0.5 ng/ml range for all assay runs.

### Placental gene expression

Placental gene expression of apelin system components and related molecules was determined by qRT-PCR. Total RNA was extracted from villous tissue using the RNeasy Plus Mini Kit (Qiagen, Germany). Concentration and purity of RNA samples were measured using a BMG LabTech FLUOstar Omega spectrophotometer. RNA concentrations ranged from 31 to 740 ng/µl with A_260_/A_280_ ratios between 1.9 and 2.1. Total RNA (0.25 µg) was reverse transcribed using the High-Capacity cDNA Reverse Transcription Kit (ThermoFisher Scientific, USA).

Expression of *APLN, APELA* and *APLNR* was determined using TaqMan probes (ThermoFisher Scientific, USA; Hs00175572_m1, Hs04274421_m1 and Hs00766613_m1). Expression of components of the renin-angiotensin system and insulin-like growth factor axis, which interact with the apelin system and are known to be important for placental function, was also measured using iTaq™ Universal SYBR Green Supermix (Bio-Rad, USA) and target specific primers. We measured expression of angiotensin II receptor type 1 (*AGTR1*), angiotensin I converting enzyme (*ACE*), angiotensin converting enzyme 2 (*ACE2*), renin receptor (*ATP6AP2*), vascular endothelial growth factor A (*VEGFA*), insulin-like growth factor 1 receptor (*IGF1R*), insulin-like growth factor 2 receptor (*IGF2R*) and insulin-like growth factor 2 (*IGF2*). Primer sequences are provided in Supplementary Table 1. We confirmed primer efficiency using a standard curve of set of serially diluted cDNA, product size using agarose gel electrophoresis and melt temperature using a melt curve. Expression of all targets was calculated using the ΔΔCt method relative to *HPRT1* and *RPS29* endogenous reference genes (Taqman probes Hs02800695_m1 and Hs03004311_m1). Relative expression values were log transformed prior to statistical analysis.

### Statistics

GraphPad Prism version 9.5.1 for Windows was used to conduct the statistical analyses. Continuous data were presented as mean ± standard deviation and categorical data were shown as percentages (%). For the plasma apelin analyses, continuous measurements were compared between pregnancy outcome groups using two-tailed, unpaired Student’s t-test. For the placental gene expression analyses, continuous measurements were compared between AGA, late-onset and term- and preterm-delivered, early-onset FGR groups by one-way ANOVA with Tukey’s post-hoc. Welch’s correction was used when variances were not homogeneous across study groups. Categorical variables were compared between groups using chi-squared test. Participants in the plasma apelin study were also separated into tertiles based on plasma apelin and the proportion of participants with each pregnancy outcome in each tertile was compared by chi-squared test. Linear regression was used to determine the relationship of plasma apelin or gene expression with clinical characteristics. Pearson’s correlation was performed to assess the interrelationship between gene expression values. In all cases, statistical significance was taken at the level P<0.05.

## Results

### Maternal plasma apelin study

#### Demographic and clinical characteristics

Women diagnosed with severe, early-onset FGR were enrolled into the study at the same gestational age, irrespective of whether they had an adverse pregnancy outcome (fetal death or livebirth <28 weeks gestational age) or a livebirth ≥28 weeks (Table 1). Maternal age and ethnicity were similar in the two groups. Participants in the adverse outcome group had a significantly higher body mass index and were more commonly diagnosed with a hypertensive disorder, compared to those in the favourable-outcome group (Table 1). None of the study participants had gestational diabetes.

**Table 1.** Demographic and clinical characteristics of participants with early-onset FGR in plasma apelin study.

|  | Early-onset FGR<br>Livebirth ≥ 28 weeks GA<br>n=31 | *n* | Early-onset FGR<br>Fetal death or livebirth < 28<br>weeks GA n=28 | *n* | P value |
| --- | --- | --- | --- | --- | --- |
| **Maternal and fetal characteristics** |  |  |  |  |  |
| Gestational age at enrolment, wk | 24.1 ± 1.5 | 31 | 23.3 ± 1.6 | 28 | P=0.054 |
| Age, years | 32.9 ± 6.2 | 30 | 33.7 ± 8.0 | 28 | P=0.677 |
| Ethnicity, *n* = Asian, Black or Mixed (%) | 11 (32.1%) | 31 | 9 (35.5%) | 28 | P=0.999 |
| Body mass index, kg/m<sup>2</sup> | 25.1 ± 5.2 | 31 | 28.4 ± 6.5 | 27 | **\*P=0.041** |
| Hypertensive disorder, *n* = diagnosed (%) | 3 (9.7%) | 31 | 10 (35.7%) | 28 | **\*P=0.026** |
| Pre-existing | 2 (6.5%) | 31 | 5 (17.9%) | 28 | P=0.240 |
| Pregnancy-induced | 1 (3.2%) | 31 | 5 (17.9%) | 28 | P=0.092 |
| Gestational diabetes mellitus, *n* = diagnosed (%) | 0 (0%) | 31 | 0 (0%) | 28 | P>0.999 |
| **Pregnancy outcome** |  |  |  |  |  |
| Incidence of IUFD, *n* (%) | 0 (0%) | 31 | 18 (64.3%) | 28 | **\*P<0.001** |
| Gestational age at delivery, wk | 33.1 ± 3.5 | 31 | 25.5 ± 1.6 | 28 | **\*P<0.001** |
| Mode of Delivery, *n* = caesarean (%) | 26 (86.7%) | 30 | 9 (34.6%) | 26 | **\*P<0.001** |
| Infant sex, *n* = female (%) | 13 (44.8%) | 29 | 6 (37.5%) | 16 | P=0.756 |
| Birth weight, g | 1412 ± 587 | 29 | 521 ± 91 | 10 | **\*P<0.001** |
| Placenta weight, g | 261 ± 91 | 8 | 97 ± 42 | 3 | **\*P=0.018** |

Intergroup comparisons were by two-tailed Student's t-test for continuous variables or Fisher's exact test for categorical variables. Statistically significant P<0.05 shown in bold. Continuous variables are shown as mean ± SD and categorical variables are shown n (%).

Intrauterine fetal death occurred in 64.3% of participants in the adverse outcome group and they had a significantly earlier average gestational age at delivery than the participants with a more favourable pregnancy outcome (Table 1). A smaller proportion of women in the adverse outcome group delivered by caesarean section. The distribution of infant sexes was similar in the two groups. Amongst the participants delivering liveborn infants, birth weight and placenta weight were significantly lower in the adverse outcome group than the group delivering ≥28 weeks gestational age (Table 1).

At enrolment into the study, ultrasound measurements of estimated fetal weight were 25% smaller in women who went on to have an adverse pregnancy outcome than in those who had a livebirth ≥28 weeks (Table 2). Estimated fetal weight z-score was also lower in the group with more adverse outcomes. Uterine artery volume flow rate was 56% lower, concomitant with higher pulsatility indices in both the uterine and umbilical arteries and lower umbilical venous blood flow in the women with adverse outcomes compared to livebirth ≥28 weeks (Table 2.) There was no significant difference in placental thickness between the groups.

**Table 2.**
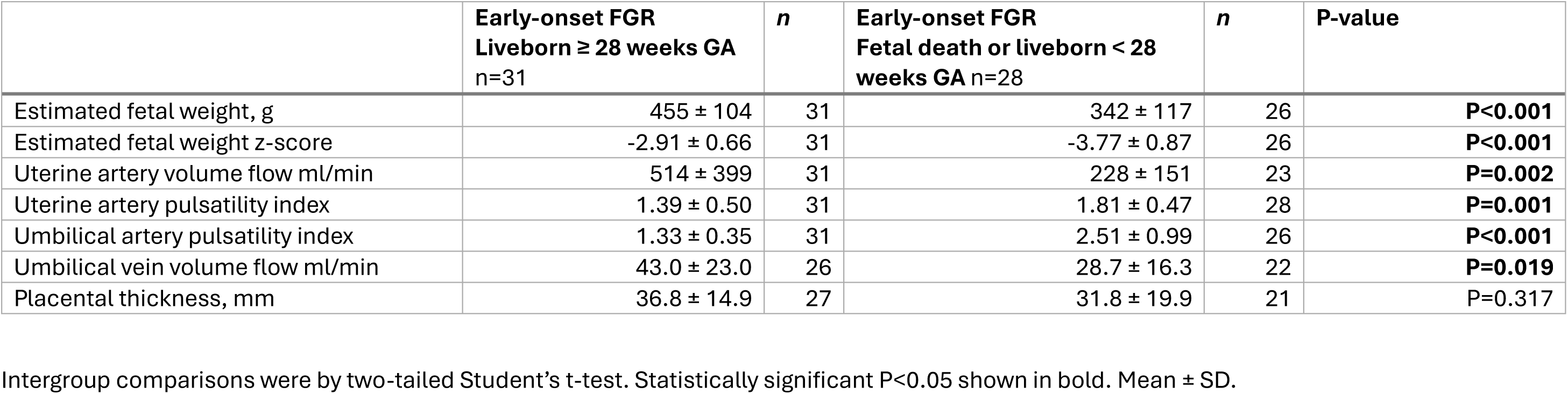
Ultrasound measurements at enrolment in participants with early onset FGR in plasma apelin study.

|  | Early-onset FGR<br>Liveborn ≥ 28 weeks GA<br>n=31 | *n* | Early-onset FGR<br>Fetal death or liveborn < 28<br>weeks GA n=28 | *n* | P-value |
| --- | --- | --- | --- | --- | --- |
| Estimated fetal weight, g | 455 ± 104 | 31 | 342 ± 117 | 26 | **P<0.001** |
| Estimated fetal weight z-score | -2.91 ± 0.66 | 31 | -3.77 ± 0.87 | 26 | **P<0.001** |
| Uterine artery volume flow ml/min | 514 ± 399 | 31 | 228 ± 151 | 23 | **P=0.002** |
| Uterine artery pulsatility index | 1.39 ± 0.50 | 31 | 1.81 ± 0.47 | 28 | **P=0.001** |
| Umbilical artery pulsatility index | 1.33 ± 0.35 | 31 | 2.51 ± 0.99 | 26 | **P<0.001** |
| Umbilical vein volume flow ml/min | 43.0 ± 23.0 | 26 | 28.7 ± 16.3 | 22 | **P=0.019** |
| Placental thickness, mm | 36.8 ± 14.9 | 27 | 31.8 ± 19.9 | 21 | P=0.317 |

Intergroup comparisons were by two-tailed Student's t-test. Statistically significant P<0.05 shown in bold. Mean ± SD.

#### Maternal plasma apelin concentrations

Women who went on to have an adverse pregnancy outcome had significantly lower plasma apelin concentrations at FGR diagnosis (15.7 ± 12.7 pg/ml) than participants with a more favourable outcome (24.6 ± 19.4 pg/ml) (Figure 1A). There was an inverse relationship between maternal plasma apelin and uterine artery pulsatility index, such that participants with higher pulsatility had lower plasma apelin concentrations (Figure 1B). Correspondingly, maternal plasma apelin correlated with uterine artery volume blood flow (Fig. 1C) but there was no significant linear relationship between maternal plasma apelin concentration and ultrasound indices of fetoplacental size or umbilical perfusion (Table 3). Plasma apelin concentrations did not significantly correlate with maternal age or body mass index (P>0.05).

**Figure 1.**
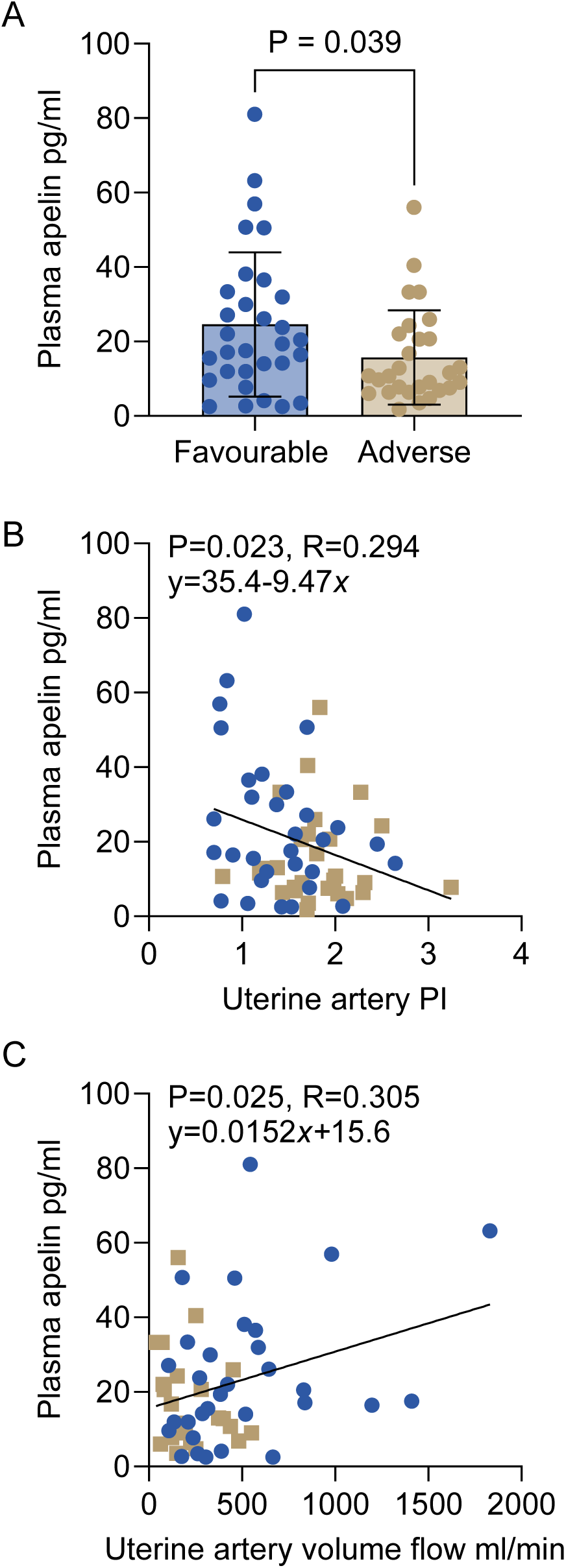
Low maternal plasma apelin concentration is associated with adverse pregnancy outcome, and uterine artery pulsatility and reduced volume flow in pregnancies complicated by early-onset FGR. (A) Maternal plasma apelin concentration at point of FGR diagnosis between 20 and 26+6 weeks gestation in women who subsequently had a livebirth ≥ 28 weeks gestational age (*n*=31) or fetal death or extremely preterm birth <28 weeks gestational age (*n*=28). Intergroup comparison by Student’s t-test with Welch’s correction. Mean ± SD, points represent individual participants. (B, C) Relationship between maternal plasma apelin concentration and uterine artery pulsatility index and volume flow rate at diagnosis with early-onset FGR (*n*=54-59). Least-squares regression line of best fit and equation shown on graph. Points represent individual participants.

**Table 3.** Linear regression relationships between ultrasound measurements and plasma apelin at early-onset FGR diagnosis.

| Variable x | P value | R value | Equation | n |
| --- | --- | --- | --- | --- |
| Estimated fetal weight | P=0.567 | R=0.077 | $y = 0.011 \text{ pg ml}^{-1} \text{ g}^{-1} x + 16.5 \text{ pg ml}^{-1}$ | 57 |
| Estimated fetal weight z-score | P=0.452 | R=0.101 | $y = 2.00 \text{ pg ml}^{-1} x + 27.4 \text{ pg ml}^{-1}$ | 57 |
| Umbilical artery pulsatility index | P=0.154 | R=0.191 | $y = -3.53 \text{ pg ml}^{-1} x + 27.0 \text{ pg ml}^{-1}$ | 57 |
| Umbilical vein volume flow ml/min | P=0.064 | R=0.270 | $y = 0.229 \text{ pg min}^{-1} x + 13.1 \text{ pg ml}^{-1}$ | 48 |
| Placental thickness | P=0.162 | R=0.205 | $y = 0.203 \text{ pg ml}^{-1} \text{ mm}^{-1} x + 13.5 \text{ pg ml}^{-1}$ | 48 |

Least-squares regression.

When the study participants were divided into tertiles based on maternal plasma apelin concentration at enrolment, there was a statistically significant overall effect of apelin tertile on pregnancy outcome (Table 4). Participants with low plasma apelin (< 10.7 pg/ml) were 1.95 times more likely to have an adverse pregnancy outcome to those with high plasma apelin (≥ 23.6 pg/ml, sensitivity 68% [confidence interval 46-85%], specificity 65% [confidence interval 43-82%]).

**Table 4.** Proportion of patients with favourable and adverse outcomes in the lower, middle, and upper tertiles of plasma apelin concentration at enrolment.

|  | Favourable (n) | Adverse (n) | P-value (chi-squared test for trend) |
| --- | --- | --- | --- |
| Lower tertile (n=20), apelin conc. <10.7 pg/ml | 7 (35%) | 13 (65%) | **P=0.036** |
| Middle tertile (n=20), apelin conc. ≥ 10.7 pg/ml and < 23.8 pg/ml | 11 (55%) | 9 (45%) |  |
| Upper tertile (n=19), apelin conc. ≥ 23.8 pg/ml | 13 (68%) | 6 (32%) |  |

### Placental apelin system expression study

#### Demographic and clinical characteristics

Women whose placentae were collected for gene expression analysis were similar in age and BMI, irrespective of whether they had early- or late-onset FGR, or delivered AGA infants (Table 5). A greater proportion of participants affected by FGR were from black, Asian or mixed ethnic backgrounds, compared to participants in the AGA group. Women diagnosed with FGR were also more commonly diagnosed with a hypertensive disorder (Table 5). The frequency of hypertensive disorders was highest in the participants with early-onset FGR delivering preterm and this was mainly due to an increased incidence of pregnancy-induced, rather than pre-existing, hypertension. There was no difference in the incidence of gestational diabetes between the groups.

**Table 5.** Demographic and clinical characteristics of participants in placental gene expression study.

|  | **AGA**<br>**Term delivery**<br>**n=15** | **n** | **Late-Onset**<br>**FGR**<br>**Term delivery**<br>**n=8** | **n** | **Early-Onset**<br>**FGR**<br>**Term delivery**<br>**n=4** | **n** | **Early-**<br>**Onset FGR**<br>**Preterm**<br>**delivery**<br>**n=16** | **n** | **P value** |
| --- | --- | --- | --- | --- | --- | --- | --- | --- | --- |
| **Maternal and fetal characteristics** |  |  |  |  |  |  |  |  |  |
| Age, years | 35.5 ± 1.5 | 15 | 32.5 ± 1.1 | 8 | 32.3 ± 3.7 | 4 | 34.8 ± 1.6 | 16 | P=0.567 |
| Ethnicity, n = Asian, Black, or Mixed (%) | 2 (13.3%) | 15 | 6 (75%) | 8 | 1 (25%) | 4 | 5 (31.3%) | 16 | **\*P=0.027** |
| Body mass index, kg/m<sup>2</sup> | 24.9 ± 1.2 | 15 | 25.4 ± 2.5 | 8 | 25.5 ± 3.2 | 4 | 28.8 ± 2.5 | 16 | P=0.509 |
| Hypertensive Disorder, n = diagnosed (%) | 0 (0%) | 15 | 1 (12.5%) | 8 | 1 (25%) | 4 | 10 (62.5%) | 16 | **\*P<0.001** |
| Pre-existing | 0 (0%) | 15 | 0 (0%) | 8 | 0 (0%) | 4 | 2 (12.5%) | 16 | P=0.593 |
| Pregnancy-induced | 0 (0%) | 15 | 1 (12.5%) | 8 | 1 (25%) | 4 | 8 (50%) | 16 | **\*P=0.009** |
| Gestational Diabetes Mellitus, n = diagnosed (%) | 0 (0%) | 15 | 1 (12.5%) | 8 | 0 (0%) | 4 | 1 (6.3%) | 16 | P=0.546 |
| **Pregnancy outcome** |  |  |  |  |  |  |  |  |  |
| Gestational age at delivery, weeks | 39.3 ± 0.2<sup>a</sup> | 15 | 37.7 ± 0.4<sup>a</sup> | 8 | 38.2 ± 0.3<sup>a</sup> | 4 | 28.9 ± 0.9<sup>b</sup> | 16 | **\*P<0.001** |
| Mode of Delivery, n = caesarean (%) | 11 (73.3%) | 15 | 7 (87.5%) | 8 | 2 (50%) | 4 | 16 (100%) | 16 | P=0.052 |
| Infant sex, n = female (%) | 5 (33.3%) | 15 | 4 (50%) | 8 | 3 (75%) | 4 | 10 (62.5%) | 16 | P=0.302 |
| Birth Weight, g | 3460.4 ± 89.1<sup>a</sup> | 15 | 2143.9 ± 169.0<sup>b</sup> | 8 | 2331.5 ± 350.7<sup>b</sup> | 4 | 884.9 ± 106.1<sup>c</sup> | 16 | **\*P<0.001** |
| Placenta weight, g† | 478.9 ± 16.6<sup>a</sup> | 9 | 321.1 ± 30.5<sup>b</sup> | 7 | 481.0 ± 103.0<sup>a</sup> | 2 | 219.3 ± 36.5<sup>b</sup> | 7 | **\*P<0.001** |

Intergroup comparisons were by one-way ANOVA for continuous variables and Chi-squared test for categorical variables. Statistically significant P<0.05 shown in bold. Different superscripts <sup>a,b,c</sup> indicate statistically different groups by Tukey's multiple comparison post-hoc test. Continuous variables are shown as mean ± SD and categorical variables are shown as n (%).

Participants in the early-onset, preterm group delivered at a significantly earlier gestational age than the other three groups, but there was no statistical difference in the rate of caesarean section or distribution of infant sexes (Table 5). Compared to AGA controls, late- and early-onset FGR were associated with a similar 33-38% reduction in infant birth weight at term. Early-onset FGR combined with preterm delivery was associated with a greater reduction in birth weight, such that the infants were 74% smaller than AGA controls and lighter than either of the other FGR groups (Table 5). By contrast, placental weight was less than AGA values in late-onset FGR and early-onset FGR, preterm groups, but not in the early-onset FGR, term group.

#### Placental gene expression

*APLN* gene expression in placental villous tissue was 63% lower in severe, early-onset FGR participants delivering preterm than in AGA controls (Fig. 2A, Tukey’s post-hoc P=0.003). *APLN* expression also tended to be lower in late-onset and early-onset FGR placentas delivered at term compared to AGA control placentas, although the difference was not statistically significant (Tukey’s post-hoc P>0.05). Expression of the apelin receptor (*APLNR*) and its other ligand elabela (*APELA*) did not differ between the groups (Fig. 2B, C).

**Figure 2.**
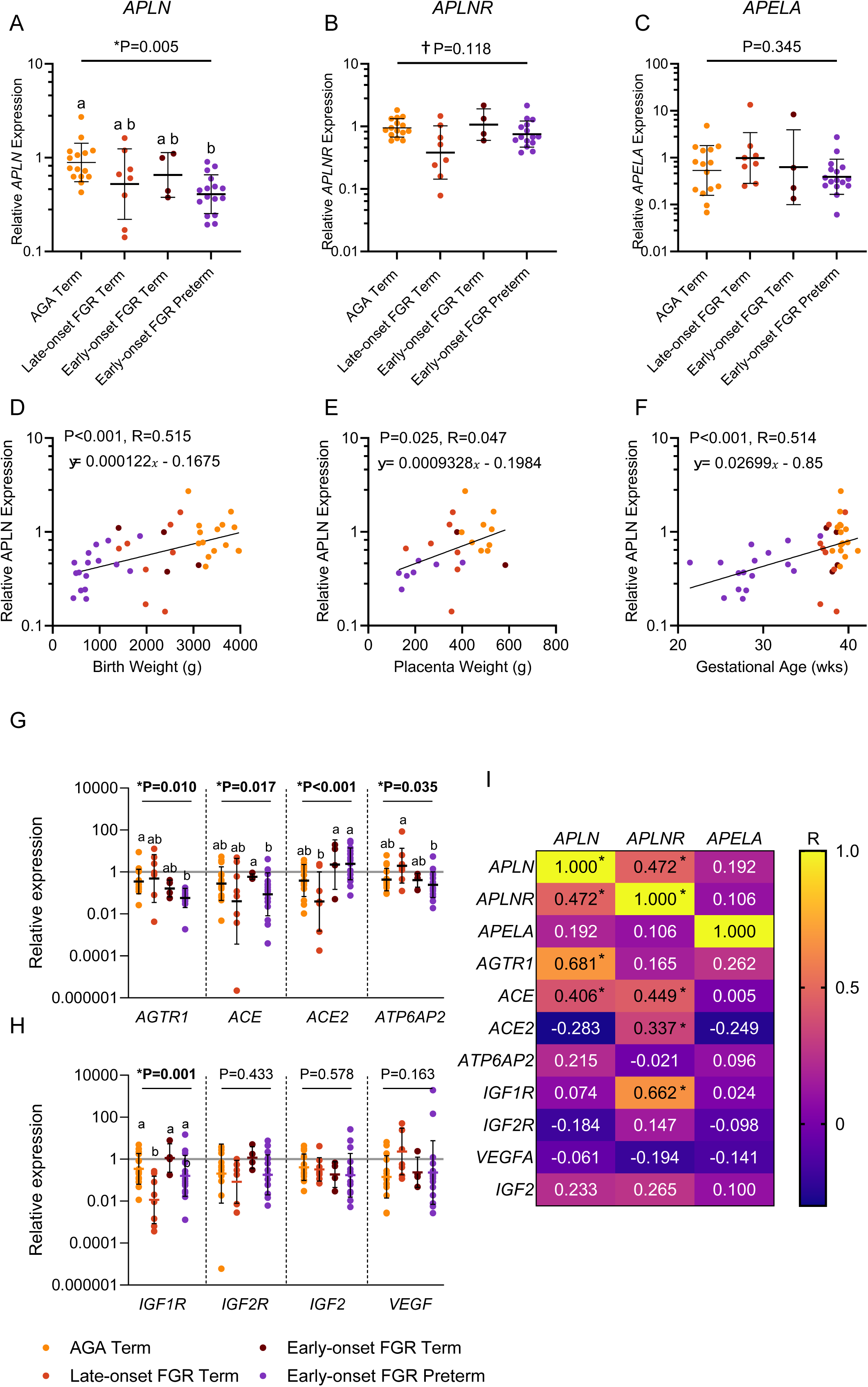
Low placental *APLN* expression in pregnancies complicated by early-onset FGR. (A-C) Expression of *APLN, APLNR* and *APELA* genes in AGA (n=15), late-onset FGR (n=8) and early-onset FGR placentas, delivered at term (n=4) or preterm (n=16). P values for one-way ANOVA shown. Different superscripts a,b represent significantly different groups by Tukey’s post-hoc test. Welch’s correction was applied for *APLNR* data due to heterogeneity of variance. Mean ± SD, points represent individual participants. (D-F) Linear relationship between placental *APLN* expression and birth weight (n=43), placenta weight (n=25) and gestational age (n=43). Regression lines and equations shown. Points represent individual participants in AGA (gold), late-onset FGR (orange), early-onset FGR born at term (brown) and preterm (purple). (G, H) Expression of renin-angiotensin system and insulin-like growth factor genes in AGA (n=15), late-onset FGR (n=8) and early-onset FGR placentas, delivered at term (n=4) or preterm (n=16). Statistics as above. Mean ± SD. (I) Correlation matrix for relationships between placental *APLN*, *APLNR* and *APELA* expression and renin-angiotensin system and insulin-like growth factor system gene expression. Values and colour scale represents Pearson correlation coefficient (R). *P<0.05 significant correlation.

When all of the groups were combined, placental *APLN* expression demonstrated a significant linear relationship with birth weight, placental weight and gestational age at delivery (Fig. 2D-F). After controlling for the effect of gestational age at delivery using multiple linear regression, there was no overall effect of FGR status on placental *APLN* expression (regression coefficients given in Table 6; main effect of FGR status P=0.356, regression fit: P=0.004, R^2^ = 0.32, degrees of freedom = 38,). Conversely, *APLNR* expression was significantly reduced in late-but not early-onset FGR placentas after controlling for gestational age (Table 6; main effect of FGR status P=0.004; regression fit: P=0.007, R^2^ = 0.31, degrees of freedom = 37).

**Table 6.** Parameters estimates for independent variables in least-squares multiple linear regression analyses of placental *APLN* and *APLNR* gene expression.

| Dependent variable | Independent variable | Regression coefficient | Standard error | P value |
| --- | --- | --- | --- | --- |
| ***APLN* expression** | Late-onset FGR | -0.192 | 0.109 | P=0.086 |
|  | Early-onset FGR, term | -0.107 | 0.138 | P=0.441 |
|  | Early-onset FGR, preterm | -0.073 | 0.188 | P=0.700 |
|  | Gestational age | 0.026 | 0.016 | P=0.120 |
| ***APLNR* expression** | Late-onset FGR | -0.363 | 0.112 | **\*P=0.003** |
|  | Early-onset FGR, term | 0.075 | 0.141 | P=0.600 |
|  | Early-onset FGR, preterm | -0.112 | 0.194 | P=0.566 |
|  | Gestational age | 0.020 | 0.017 | P=0.225 |

In the renin-angiotensin system, placental *AGTR1* expression was lower in early-onset FGR participants delivering preterm, but not in either of the other FGR groups, compared to AGA controls (Fig. 2G). Placental *ACE*, *ACE2* and *ATP6AP2* expression did not differ from AGA control values in any of the FGR groups, although there were significant intergroup differences between the FGR subtypes. Placental *ACE* was downregulated in women with early-onset FGR delivering preterm, whilst *ACE2* was upregulated and *ATP6AP2* tended to be downregulated in early-compared to late-onset FGR placentas (Fig 2G). *IGF1R* expression was downregulated in late-but not early-onset FGR placentas compared to AGA controls but there was no effect of FGR status on placental *IGF2R, IGF2* or *VEGF* expression (Fig 2H).

When all of the groups were combined, placental *APLN* expression correlated with *APLNR, AGTR1* and *ACE* expression (Fig 2I). *APLNR* expression also correlated with *ACE*, as well as *ACE2* and *IGF1R* expression.

## DISCUSSION

In this study we determined systemic concentrations and placental expression of apelin in pregnant women diagnosed with severe, early-onset FGR. Low plasma apelin during the second trimester at FGR diagnosis was associated with subsequent intrauterine fetal death or extreme preterm delivery. Low maternal plasma apelin was also associated with higher uterine artery pulsatility, a measure of uteroplacental blood flow resistance, and lower uterine artery volume blood flow. Placental gene expression of *APLN*, but not *APLNR*, was downregulated in early-onset FGR and correlated with both fetoplacental weight and gestational age. By contrast, late-onset FGR was associated with lower placental expression of the receptor apelin *APLNR*, but not *APLN,* when gestational age was accounted for. There were positive correlations between placental expression of the apelin system genes and components of the renin-angiotensin system and insulin-like growth factor system. Taken together, the findings indicate that apelin signalling plays an important role in placental function and fetal growth and that apelin deficiency could underpin adverse pregnancy outcome in pregnancies complicated by FGR.

The plasma apelin concentrations measured in our study were on average lower than the range of values previously reported in pregnant women [14, 26]. We used solid-phase extraction to remove interfering proteins from the plasma samples prior to the apelin immunoassay, whereas the previous studies did not include this step. Peptide extraction has been shown to permit reproducible and sensitive measurement of [Pyr]^1^-apelin-13 by immunoassay [25] and is necessary for accurate measurement of plasma elabela during pregnancy [27]. Therefore, our apelin concentration measurements were more similar to previously published values in healthy nonpregnant individuals, determined with the extraction step [28]. Our results are also consistent with lower serum apelin levels in women with FGR-complicated pregnancies, when compared to uncomplicated pregnancies [14]. Our data additionally show that maternal plasma apelin concentration is related to FGR severity, because fetuses in the group with the most adverse pregnancy outcomes were already smaller at the point of diagnosis.

The link between low maternal plasma apelin and adverse pregnancy outcome could be attributed to maternal disease. Participants in the study group with the worst pregnancy outcomes had higher average body mass index and a greater rate of hypertension than the participants with better outcomes. Pre-pregnancy obesity and hypertension are both associated with increased risk of preterm birth [29, 30]. Apelin is highly expressed in adipocytes and endothelial cells [31], so the circulating peptide could originate from those cell types and indicate alterations in their number or function related to cardiometabolic disease. However, plasma apelin *increases* in non-pregnant people with obesity [32] and was not affected by obesity in our previous study of pregnant women [33]. Conversely, plasma apelin decreases in patients with primary hypertension or pregnancy-induced pre-eclampsia, similar to the association with adverse pregnancy outcome and concomitant with decreased apelin expression in the endothelium [34, 35]. Therefore, lower apelin levels in the adverse pregnancy outcome group are more likely to be a marker of hypertension and maternal vascular disease leading to preterm delivery. We were not able to measure maternal plasma apelin in the first trimester so it remains unclear whether maternal apelin levels independently relate to pre-existing maternal disease, independent of FGR diagnosis.

Lower apelin levels in the adverse outcome group may alternatively be linked to poor placental function. Apelin expression in the human term placenta is higher than in adipose tissue [11] and maternal serum apelin levels correlate with placental apelin expression in FGR [14]. The inverse relationship between low circulating apelin and high uteroplacental resistance suggests that impaired trophoblast cell invasion could explain the low apelin levels. Defective trophoblast invasion into the uterine decidua is a hallmark of FGR and associated with Doppler indices of high uterine artery resistance [36]. Invasive extravillous trophoblast cells express apelin [37] and placental explants secrete apelin *in vitro* [15]. Rat placental tissue cultured *ex vivo* also secretes apelin into the medium [17]. Therefore, low plasma apelin levels in the participants who went on to have an intrauterine death or severe preterm delivery may be due to defective function of placental trophoblast cells.

Our finding of lower *APLN* expression specifically in early-onset FGR compared to AGA placentas reflects previous data showing reduced placental apelin protein abundance in a mixed FGR population defined based on birth weight [14]. By contrast with the previous study, we demonstrated *APLN* downregulation at the transcript level in FGR placentas. This may be because we analysed early and late-onset FGR placentas separately and used a slightly larger sample size. Although we did not measure cell-specific expression, the previous publication demonstrates that the apelin protein is downregulated in FGR in the syncytialised trophoblast cells of the placental epithelium [38]. Lower placental *APLN* expression in early-onset FGR participants delivering preterm could be explained by the confounding effect of gestational age, which was less than in the AGA control group and also positively correlated with *APLN* expression. Our study was limited by the lack of a preterm, AGA group of participants. However, apelin abundance does not increase with gestational age in either humans or rodents when it is measured longitudinally [17, 38], so it is more likely that downregulation here is associated with FGR *per se*. Our results are also in line with decreased expression of apelin and *APLNR* in placentas from patients with pre-eclampsia, another pregnancy complication characterised by placental insufficiency [13, 15]. Overall, they support the concept that early-onset FGR is associated with placental apelin deficiency.

Our finding that placental *APLNR* expression is reduced specifically in late-onset FGR is novel to the best of our knowledge. Undernutrition in pregnant rats reduces placental *Aplnr* in line with reduced fetal weight [39] and pre-eclampsia is associated with reduced placental *APLNR* in humans [40]. Similar to *APLN*, *APLNR* is localised to the syncytiotrophoblast and endothelial cells in the human placenta [40]. Reduced *APLNR* expression could reflect defective function of either of these cell types and points to different pathophysiological mechanisms underpinning late- and early-onset FGR. We did not find any effect of FGR on placental expression of *APELA,* consistent with previous studies measuring both mRNA and protein expression of elabela in placentas affected by pre-eclampsia [37, 41].

The correlations in placental expression of apelin system and renin-angiotensin system components are consistent with their antagonistic actions and the established spatial colocalization of their main receptors. Angiotensin II increases uteroplacental vascular resistance by constricting arteries on both the maternal and fetal side of the placenta via AGTR1 [42, 43]. It may also affect trophoblast invasion because *AGTR1* is expressed in extravillous trophoblast cells [44]. Angiotensin II downregulates apelin expression including in the placenta [15, 45]. Apelin promotes angiotensin II breakdown by upregulating ACE2 [46] and causes antagonistic vasodilatation. The strong correlation between *APLN* and *AGTR1* expression in our study was driven by downregulated *AGTR1* in the preterm, early-onset FGR group, consistent with previous data showing that *AGTR1* is reduced in FGR placental endothelium and trophoblast [47, 48]. Combined with the tendency for lower expression of the activating components of the renin-angiotensin system, *ACE* and *ATP6AP2*, and higher inactivating *ACE2*, this suggests dampened local production of angiotensin II most likely due to negative feedback from increased fetal angiotensin II concentrations in FGR [49]. Our data may therefore also suggest that placental *APLN* is suppressed by increased fetal systemic angiotensin II availability in early-onset FGR.

The strong positive correlation between *APLNR* and *IGF1R* expression appeared to be driven by decreased *IGF1R* expression in late-onset FGR placentas. Although we were only able to measure mRNA expression, IGF1R protein abundance is commonly reduced in FGR human placentas [50] and in pregnant rats following uterine artery ligation [51]. Genetic studies consistently demonstrate that *IGF1R* mutations and polymorphisms contribute to FGR and normal birth weight variability [52, 53]. Loss-of-function mutations in *Igf1r* reduce fetal weight in mice [54]. *IGF1R* is expressed in all placental cell types and activation by either IGF1 or IGF2 increases growth and nutrient uptake in trophoblast cells [55] and transplacental nutrient transport in pregnant animals [56]. APLNR and IGF1R share common downstream signalling mechanisms, both activating kinase signalling through protein kinase B and mechanistic target of rapamycin. There appears to be crosstalk between the two receptors in cancer cells, but little other information exists on their relationship [57]. Our data suggest that *IGF1R* and *APLNR* expression are commonly regulated and that altered APLNR signalling could be a cause or consequence of altered IGF1R signalling in FGR placentas.

In conclusion, our results agree with the original hypothesis that extreme preterm delivery and intrauterine fetal death are associated with low maternal plasma apelin at the point of diagnosis with fetal growth restriction. They also confirm that placental *APLN* gene expression is lower in early-onset FGR than in normally-grown control pregnancies. The findings improve our understanding of the distinct pathophysiological processes contributing to early- and late-onset FGR. Since apelin deficiency preceded adverse pregnancy outcome, the findings may mean that apelin is needed for normal fetal growth and maintenance of pregnancy, for example to promote placental function or mediate cardiovascular adaptations. Apelin deficiency could therefore mechanistically underpin impaired placental function and fetal growth. Certainly, apelin receptor agonists enhance trophoblast cell invasion and nutrient uptake *in vitro* [20, 33, 58, 59] and stimulate placental amino acid transporter expression and glucose transfer *in vivo* [39, 60]. We speculate that measuring or augmenting apelin availability or apelin receptor signalling could be an effective intervention to prognose or treat severe, early-onset FGR.

## Funding

The research leading to these results received funding from The Royal Society (to ORV); the Elizabeth Garrett Anderson Hospital Charity (to ORV); the European Union Seventh Framework Programme (FP7/2007-2013) under grant agreement no. 305823; the Rosetrees Trust; and the Mitchell Charitable Trust in memory of Shoshana Mitchell Glynn. This research has also been supported by the National Institute for Health Research University College London Hospitals Biomedical Research Centre (to RS, ALD).

---

**Supplementary Table 1.**
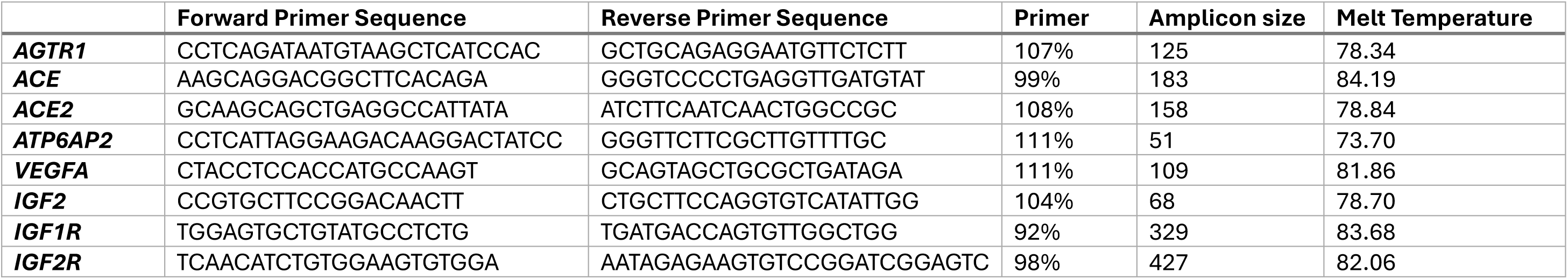
Sequences and optimisation details for qRT-PCR primers.

## Notes

### Competing Interest Statement

The authors have declared no competing interest.

